# Microscale Spatial Dysbiosis in Oral biofilms Associated with Disease

**DOI:** 10.1101/2024.07.24.604873

**Authors:** Benjamin Grodner, David T. Wu, Sumin Hahm, Lena Takayasu, Natalie Wen, David M. Kim, Chia-Yu Chen, Iwijn De Vlaminck

## Abstract

Microbiome dysbiosis has largely been defined using compositional analysis of metagenomic sequencing data; however, differences in the spatial arrangement of bacteria between healthy and diseased microbiomes remain largely unexplored. In this study, we measured the spatial arrangement of bacteria in dental implant biofilms from patients with healthy implants, peri-implant mucositis, or peri-implantitis, an oral microbiome-associated inflammatory disease. We discovered that peri-implant biofilms from patients with mild forms of the disease were characterized by large single-genus patches of bacteria, while biofilms from healthy sites were more complex, mixed structures. Based on these findings, we propose a model of peri-implant dysbiosis where changes in biofilm spatial architecture allow the colonization of new community members. This model indicates that spatial structure could be used as a potential biomarker for community stability and has implications in diagnosis and treatment of peri-implant diseases. These results enhance our understanding of peri-implant disease pathogenesis and may be broadly relevant for spatially structured microbiomes.

## Introduction

Microbial biofilm-associated diseases around dental implants represent a significant medical and socioeconomic burden.^1–3^ Peri-implant diseases are difficult to treat despite existing technologies, and have high recurrence following therapy.^4^ Studies of peri-implant biofilms have revealed differences in bacterial composition between peri-implant health and peri-implant diseases.^5–9^ In the microbiome field, this shift to a disease-state community composition is termed “dysbiosis”.^10–12^ Yet defining dysbiosis as such is incomplete; surface-associated microbiomes such as peri-implant biofilms are taxonomically diverse and spatially structured.^13–19^ The recent development of highly multiplexed fluorescence in situ hybridization (FISH) enables the exploration of the spatial architecture of microbiomes.^13,20–22^ In this study, we used high phylogenetic resolution fluorescence in situ hybridization (HiPR-FISH) to map the micron scale spatial structure of peri-implant bacteria in biofilms collected from patients with healthy peri-implant tissue, peri-implant mucositis, and peri-implantitis.^1,23,24^ We observed different spatial patterns between peri-implant health, peri-implant mucositis, and peri-implantitis biofilms, suggesting that changes in biofilm architecture are a feature of dysbiosis.

Since the spatial data from peri-implant biofilms is highly variable, the visually apparent patterns in the data are challenging to quantify. To establish metrics for biofilm spatial structure, we turned to examples from macroscale ecology, specifically power laws and species-area curves.^25,26^ Power laws have been used to describe bacterial clustering in model systems.^27–29^ In peri-implant biofilms, we found that the bacterial density follows a power law and that these can be used to differentiate visually distinct spatial distributions. The species-area relationship is widely used for ecological applications and can be adapted to include both species richness and relative species abundance.^30^ We used the species-area relationship as a metric for spatial heterogeneity of bacterial taxa across length scales. This metric quantifies the patterns we observed in the data and can be used to compare biofilm spatial patterns across many heterogeneous samples, enabling larger clinical studies of spatial patterns in microbiome dysbiosis.

## Results

We recruited patients with dental implants at the Harvard School of Dental Medicine and collected samples by scraping with curettes and storing subgingival peri-implant biofilm. We imaged samples from 25 donors: 6 were diagnosed with peri-implant health, 7 diagnosed with peri-implant mucositis, 7 diagnosed with mild peri-implantitis, and 5 diagnosed with moderate/severe peri-implantitis. Sample sites with mild or moderate/severe peri-implantitis had peri-implant pocket depth greater than 5 mm with bleeding on probing and/or suppuration. Mild peri-implantitis sites showed radiographic evidence of less than 30% bone loss around the implant, while moderate/severe peri-implantitis sites showed bone loss greater than 30-50% around the implant.

To stain the samples, we selected a panel of HiPR-FISH probes targeting 18 abundant and prevalent genera in the Human Oral Microbiome Database and assigned a combinatorial fluorescent barcode to each genus.^14^ Using confocal microscopy with spectral detection, we imaged the samples and collected tile scans with multiple adjacent fields of view (FOV). In total, we captured 63 tile scans comprised of 418 FOV. Using automated image segmentation, we identified 4,178,996 labeled bacterial cells, which we then classified using their fluorescent spectra and generated spatial maps of genus identity (**Fig. 1a-e**). We then visually inspected these genus-classified bacterial maps to identify differential spatial patterns between clinical diagnosis groups.

**Figure 1.**
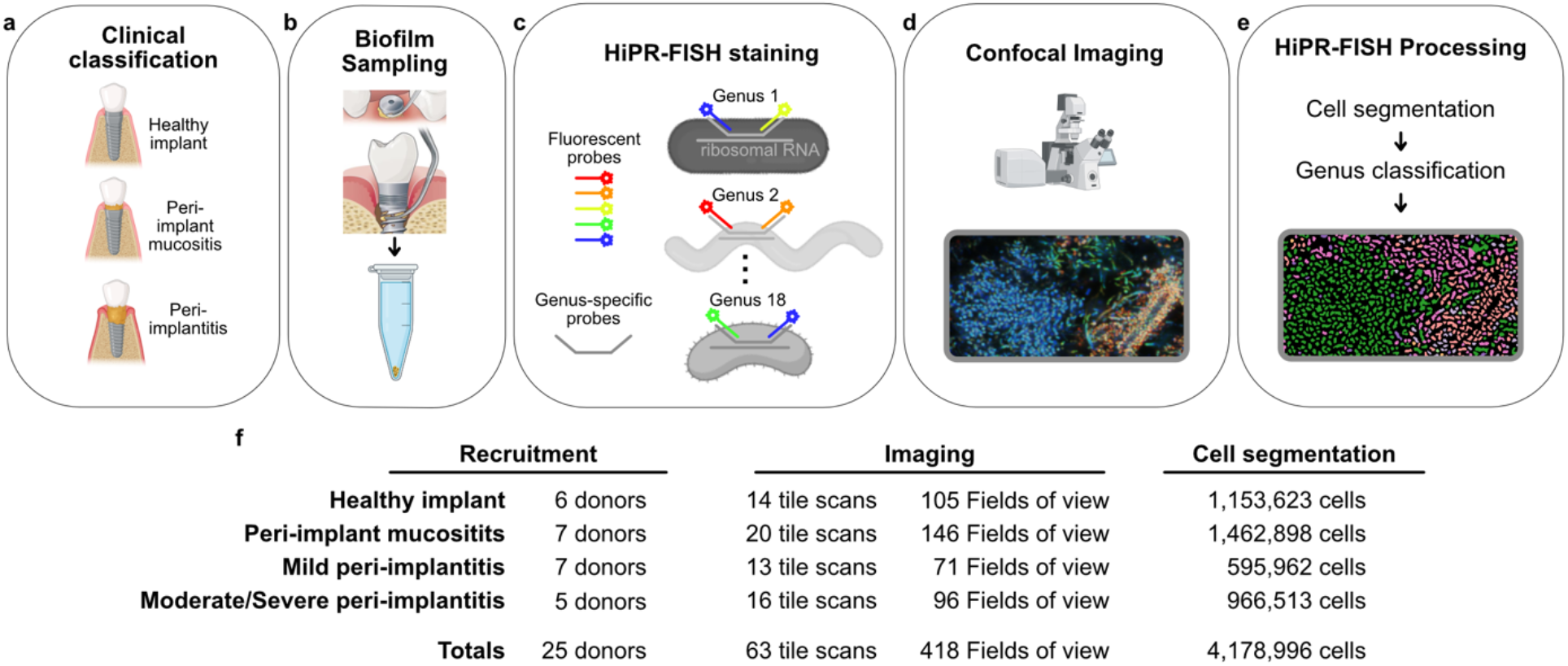
Data Collection Workflow: Patients were diagnosed with peri-implant health, peri-implant mucositis, and peri-implantitis (**a**), and biofilm samples were collected around dental implants (**b**). Samples were stained with HiPR-FISH fluorescent genus-specific probes (**c**) and imaged on a confocal microscope (**d**). The data was processed using HiPR-FISH cell segmentation and genus classification pipelines (**e**). **f** Breakdown of values for sampling, imaging, image processing, and image analysis across clinical classifications, which are defined in bold in the leftmost column. Underlined column labels show the processing workflow from left to right.

We discovered that mild peri-implantitis biofilms were dominated by large, 50-100 μm diameter single-genus regions, while the largest regions in peri-implant health biofilms were 5-25 μm (**Fig. 2**). Peri-implant mucositis and mild peri-implantitis biofilms commonly exhibited sharp borders between single-genus regions, while these borders were more mixed in peri-implant health biofilms. Moderate/severe peri-implantitis biofilms exhibited both large, single-genus patches and regions of well-mixed taxa on the scale of 25μm.

**Figure 2.**
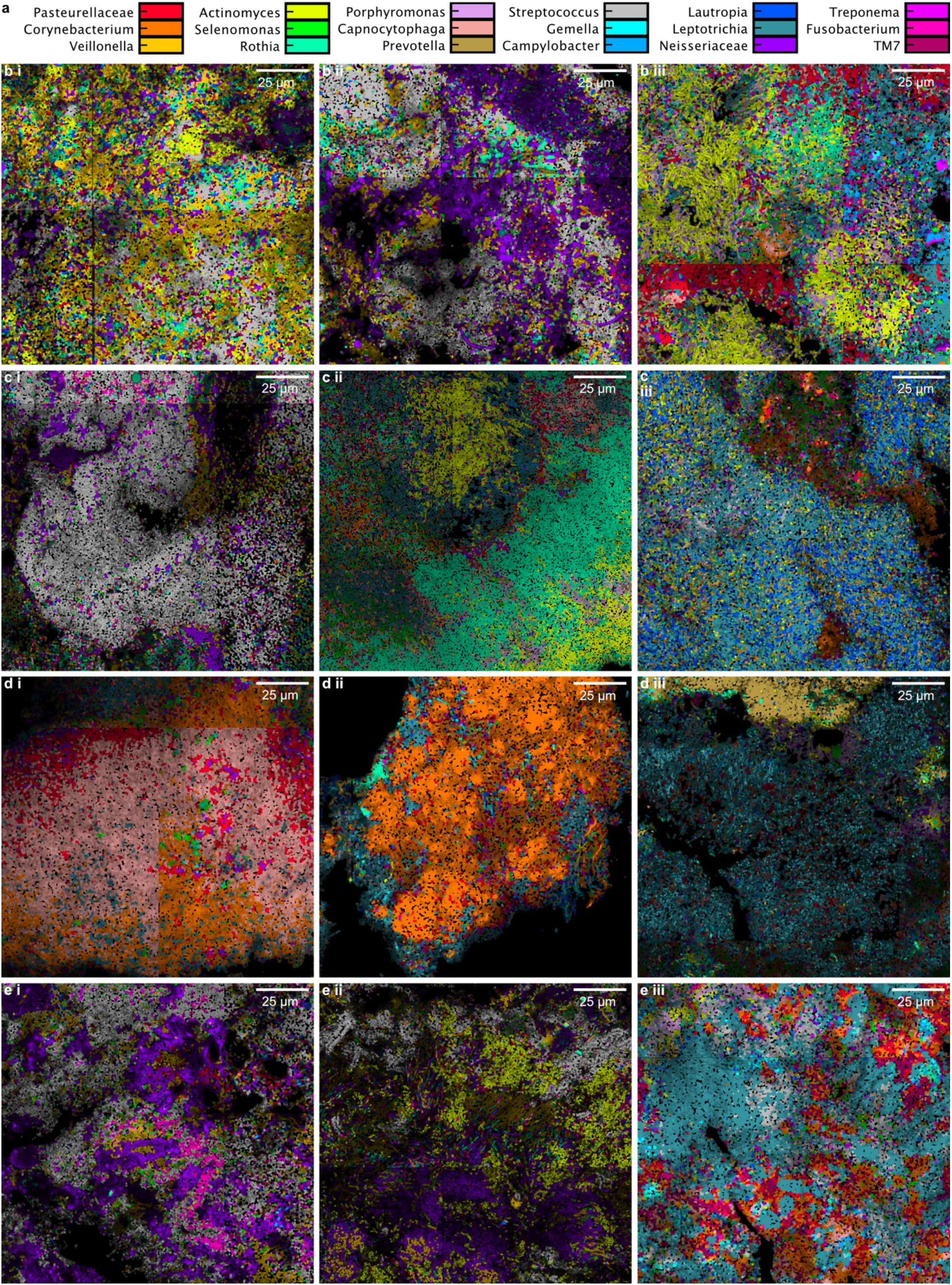
Example processed images: **a** Color legend for classified cells. **b** Segmented and classified tile scans of biofilms from peri-implant health. **c-e** Same as **b** but for peri-implant mucositis, mild peri-implantitis, and moderate/severe peri-implantitis, respectively.

To quantify these observations, we first measured the power spectrum of genus density.^31,32^ Like many other biological systems, we found that genus density follows a power law over the scales considered.^26^ We found that power law exponents could be used as metrics to differentiate visually distinct patterns between genera (**Extended data Fig. 1**). We next sought to describe the overall multi-genus spatial patterns we observed in each tile scan. We found no significant differences between groups in mean alpha diversity at a range of scales (**Extended data Fig. 2**). We used a box-counting method to measure the scaling of local genus-level diversity (**Fig. 3a**).^30^ The slope of Simpson diversity versus area on a log-log scale describes the homogeneity of the sample, where a slope of 2 indicates perfectly mixed genera and a slope of 1 indicates the perfect separation of genera.^33^ We used the Simpson diversity-area relationship as a metric to summarize the group differences between peri-implant health, peri-implant mucositis, mild peri-implantitis, and moderate/severe peri-implantitis (**Fig. 3b**). We showed that healthy peri-implantitis biofilms were more well mixed than mild peri-implantitis and moderate/severe peri-implantitis biofilms (Kruskal-Wallis H-test p = 0.018, Dunn’s test p = 0.002 and p = 0.018 respectively, **Figure 3b**). This analysis enables to quantify visually apparent trends in multi-genus spatial structures across heterogeneous peri-implant biofilm samples.

**Figure 3.**
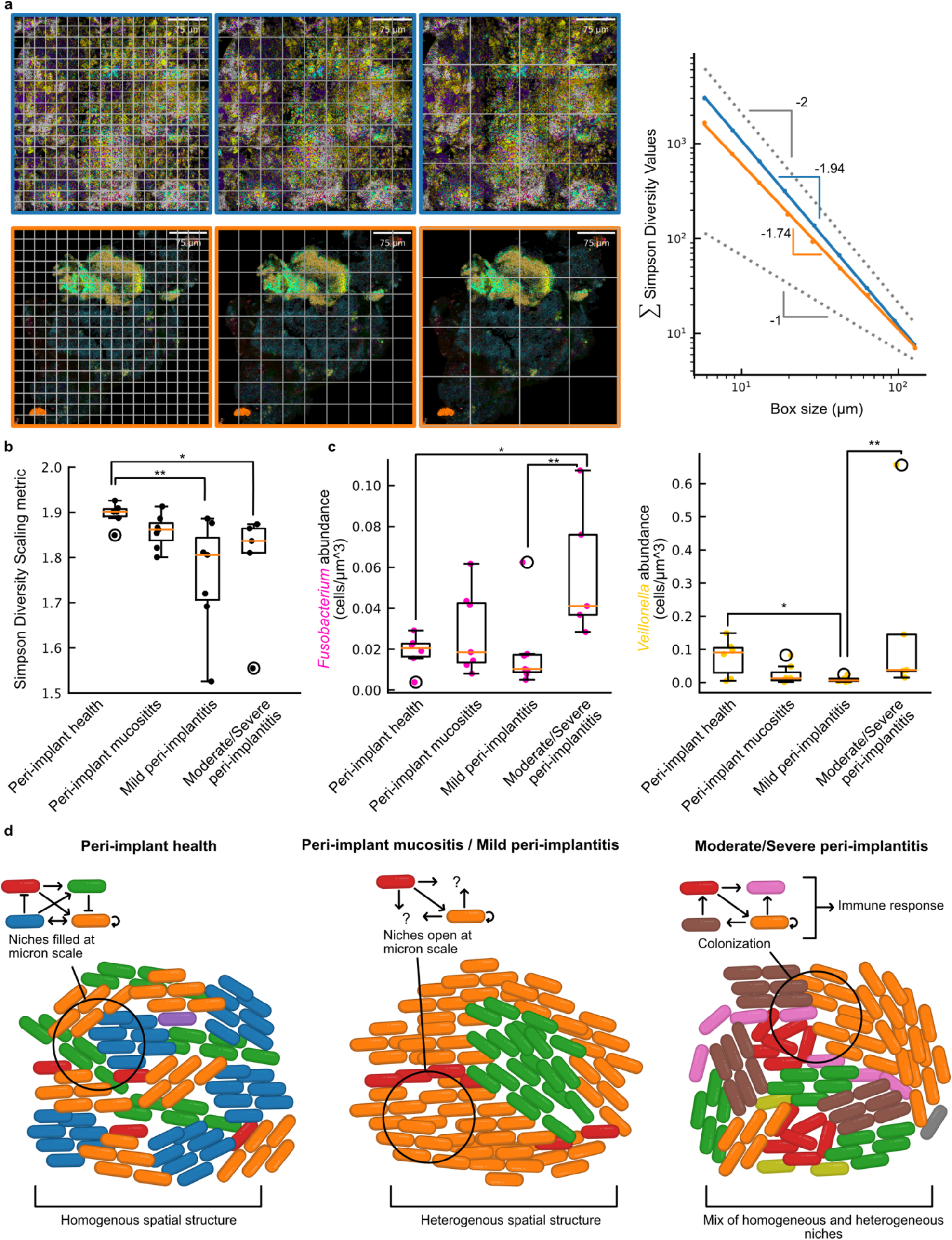
Spatial structure analysis: **a** *Left:* Diagrams illustrating the implementation of increasing box size used to measure the scaling of local diversity on two images. The Simpson diversity value is calculated for each box. *Right:* Plot of summed Simpson diversity values versus box size and resulting linear regression on a log-log scale for the *upper* image (blue points and upper solid blue line) and the *lower* image (orange points and lower blue line). The upper dashed gray line illustrates a slope of -2 and the lower dashed gray line illustrates a slope of -1. The slope of the regression is multiplied by -1 and used as the Simpson diversity scaling metric. **b** The Simpson diversity scaling metric for all tile scans grouped by clinical status. **c** *Left*: Absolute abundance of *Fusobacterium* measured as cells/μm^3 in each donor grouped by clinical status. *Right:* The same plot for *Veillonella*. **d** Proposed model of dysbiosis where peri-implant health biofilms are stable because niches are filled at the small scale. In mucositis, the biofilm is destabilized and large low diversity patches regions with unfilled niches. In peri-implantitis, new colonizers result in new interactions and the new community aggravates the host immune system. In **b** and **c**, pairwise comparison of groups was performed using Dunn’s test. The * symbol indicates p-value ≤ 0.05, the ** symbol indicates p-value ≤ 0.01, and a lack of symbol indicates p-value > 0.05. The bounds of the boxes show the first quartile to the third quartile, the center shows the median, and the whiskers show the farthest data point lying within 1.5x the inter-quartile range.

To evaluate differences in composition between groups, we quantified the absolute abundances of genera in each tile scan, counting the number of cells and dividing by the imaging volume of the confocal optical sectioning. We found that *Fusobacterium* abundance was increased in moderate/severe peri-implantitis biofilms compared to peri-implant health and mild peri-implantitis biofilms (Kruskal-Wallis H-test p = 0.049, Dunn’s test p = 0.045 and p = 0.006 respectively, **Figure 3c**). We also found that *Veillonella* abundance was decreased in mild peri-implantitis biofilms compared to peri-implant health and moderate-severe peri-implantitis biofilms (Kruskal-Wallis H-test p = 0.024, Dunn’s test p = 0.022 and p = 0.007 respectively, **Figure 3c**). Other studies have found that *Fusobacterium* is associated with peri-implantitis and *Veillonella* is associated with peri-implant health.^34,35^ Our findings provide direct evidence for the previous sequencing-based association of *Fusobacterium* and *Veillonella* abundance with moderate/severe peri-implantitis and peri-implant health respectively.

## Discussion

In this study, we used bacterial spatial mapping to reveal differences in the spatial structure of peri-implant biofilms between healthy and disease states. To our knowledge, this is the first time spatial measurement at single-cell resolution has been used to compare health- and disease-associated biofilms. We observed that peri-implant mucositis and mild peri-implantitis biofilms were characterized by large single-genus patches, while healthy biofilms had smaller mixed patches, and moderate/severe peri-implantitis biofilms had a combination of the two patterns. Previous compositional comparisons have shown that in peri-implant mucositis, there is a distinct microbial composition from both healthy sites and peri-implantitis sites, representing an intermediate condition.^7,36^ Our observation of plaque spatial structure supports the concept of mucositis as an intermediate stage between healthy and severe peri-implantitis biofilms.

To enable comparisons across many samples, we evaluated metrics of spatial structure from ecology. We found that the genus-level abundances in plaque biofilms are self-similar over the measured scales, following a power law for the power spectrum of cell density. We also found that the power law relationship between Simpson diversity and area could be used as a metric for spatial homogeneity and could differentiate the clinical groups. This metric generalizes our qualitative observations and will enable quantitative comparison of many samples in clinical studies. We also demonstrated the utility of imaging as a direct measure of absolute genus abundance in units of cells per unit volume, validating previously reported associations of genus-level abundance with peri-implant health and moderate/severe peri-implantitis.

Based on our observations, we propose a model of peri-implant dysbiosis that incorporates changes in biofilm spatial structure (**Fig. 3d**). In this model, peri-implant health biofilms are stable and resistant to pathogen colonization because they form spatial structures with high local diversity such that functional niches are filled at the 10-100 μm scale.^17^ In peri-implant mucositis, the bacterial ecosystem becomes unstable: there are blooms that form large patches, and these patches open niches for colonization by new community members or an opportunistic existing community member since functional roles are missing. Moderate/severe peri-implantitis biofilms have established new stable micron-scale spatial structures where the set of interactions antagonizes the immune system, for example the loss of *Veillonella* and the increased abundance of *Fusobacterium* as an opportunistic pathogen.^35^ We suggest that microbiomes are resilient in health and dysbiosis. In moderate/severe peri-implantitis, where the dysbiotic microbiome has established a stable community with high local diversity, the resilience of this dysbiotic state makes it much more difficult to treat. This could explain the unpredictable peri-implantitis treatment often observed in the clinic.^1^

Our model of peri-implant dysbiosis introduces immediate questions for further investigation and has implications for the treatment of peri-implant disease. The factors destabilizing a peri-implant health biofilm and causing a shift to a peri-implant mucositis and mild peri-implantitis biofilm remain unclear. If the shift to peri-implant mucositis-like biofilm structures precedes the development of clinical symptoms, the taxonomic spatial structure could be used as a biomarker for microbiome instability and the prediction of dysbiosis. Dysbiotic biofilms in moderate/severe peri-implantitis could be perturbed to an unstable state, as quantified by spatial metrics, such that new colonization becomes possible and then seeded with a desirable community as in fecal microbiome transplants.^37^ For patients with biofilms already in unstable intermediate states such as peri-implant mucositis and mild peri-implantitis, microbiome transplant methods could be used as a prophylactic. Finally, this model of dysbiosis may be generalizable to periodontal biofilms as well as other spatially structured communities.

## Methods

### Patient recruitment

Patients were included in recruitment by age greater than 18 years and by the presence of osseointegrated dental implant in peri-implant health or disease states. Patients were excluded from recruitment for uncontrolled medical conditions, type 1 and 2 diabetes mellitus, untreated periodontal conditions, use of systemic antibiotics, and smoking more than 20 cigarettes a day.

### Sample collection

Samples of human subgingival plaque biofilm are collected from patients with peri-implant health, peri-implant mucositis, or peri-implantitis. The study was approved by the Harvard Faculty of Medicine Office of Regulatory Affairs and Research Compliance and Institutional Review Board (IRB21-0662). Informed consent is obtained from all patients. The subgingival plaque was collected using a plastic curette, gently deposited into 1 ml of 70% ethanol, and stored at -20 °C until use. Plaque samples from 4 sites around one dental implant per patient were collected (Mesio-buccal, disto-buccal, mesio-lingual and disto-lingual). Full mouth plaque score and bleeding score were measured at the time of plaque collection and probing pocket depth at 6 sites around the implant and soft tissue recession around the dental implant. A periapical radiograph was also performed.

### Diagnosis

Peri-implantitis was diagnosed based on probing pocket depth greater than 5 mm, peri-implant sites with bleeding on probing, and radiographic evidence of greater than 2 mm of marginal bone loss or exposure of one implant thread compared with the bone level on a previous radiograph. Mild peri-implantitis was defined as radiographic evidence of bone loss of less than 30% around the implant, greater than 5 mm probing pocket depth, and bleeding on probing. Moderate to severe peri-implantitis was defined as bone loss greater than 30-50% around the implant, greater than 5 mm probing pocket depth, and bleeding on probing.^38^ Peri-implant mucositis was diagnosed by visual assessment of tissues as red, swollen, and soft compared with baseline as well as the presence of bleeding or suppuration on probing and an increase in probing depths compared to baseline. Peri-implant mucositis was further defined by the absence of bone loss beyond crestal bone level changes resulting from the initial remodeling.^39^

### Calibration

Clinical measurements were performed by one of two examiners at the clinical test site. Training and calibration sessions were conducted with the PI (D.M.K) until a ± of 1-mm or less in the IPD was achieved. A Kappa-statistics of 0.8 or greater was required.

### Fluorescent readout probes

Fluorophore conjugated oligonucleotide readout probes were purchased from Integrated DNA Technologies (IDT) and Biosynthesis. (Table reference)

### Ribsomoal RNA encoding probes

Probes were selected from a previous study targeting abundant and prevalent taxa in dental plaque at the genus level ^14^. Barcodes were assigned to genus-level probes by appending flanking sequences (the reverse complement of a fluorescent probe) to the 5 prime and 3 prime end of probe. Barcodes were selected such that the hamming distance between barcodes was maximized. Probes were purchased form IDT.

### HiPR-FISH staining

HiPR-FISH staining was conducted as previously described.^13^ In short, a plaque sample was deposited on a microscope slide and heated briefly to attach the sample to the surface. The sample was then briefly fixed using formaldehyde and the cell walls of the bacteria were permeabilized with lysozyme. The sample was covered in a hybridization buffer containing a high concentration of the complex oligo pool and incubated overnight. The sample was then washed to remove excess probe, covered in a hybridization buffer containing fluorescently labeled readout probes, and incubated for two hours. The sample was washed a final time, then embedded in an antifade mountant and sandwiched between a microscope coverglass.

### Imaging

Spectral images were recorded on an inverted Zeiss 880 confocal microscope equipped with a 32-anode spectral detector, a Plan-Apochromat 63X/1.40 oil objective and excitation lasers at 405 nm, 488 nm, 514 nm, 561 nm, 633 nm using acquisition settings as previously described.^21^ X-y locations were selected manually by searching the slide for intact plaque material. The z-locations for each x-y location were selected by visually assessing the highest signal intensity in the z direction. Tile scan images of varying sizes were captured for each location at three (488nm, 514nm, 561nm) or four (488nm, 514nm, 561nm, and 633nm) laser wavelengths depending on the barcoding scheme. Tile scan size was determined manually by evaluating the size of the sample fragment at the location. Where samples resulted in a large mass of plaque, tile scans were collected at three locations. For samples with less than three pieces of intact plaque, tile scans were collected from all intact regions.

### Data exclusion

Each spectral image was summed along the spectral dimension and then the summed images from the first three lasers (488nm, 514nm, 561) were stacked into an (x,y,3) array and displayed as an RGB image. Tile scans were visually inspected excluded from further analysis based on visual assessment of the absence of bacteria. Examples of excluded versus included images are shown in **Extended Data Fig. 3**. All excluded images are indicated in the “Excluded” column in **Table 5**.

### Image Processing

Images were processed using a custom Python pipeline.^40^ For a given tile, the captured images from different lasers were registered to each other in the X-Y direction to correct for shifts. The registered images from different lasers were concatenated along the spectral dimension. The maximum intensity projection along the spectral dimension was then used for automated segmentation of single cells. In brief, cell edges were highlighted using a custom algorithm called local neighborhood enhancement, the background was subtracted, and the resulting image was segmented using a watershed algorithm. Tiles were then stitched into a complete image. Cell locations were determined using the centroid of the segmentation objects. To determine the fluorescent barcode of each cell, we first calculated the average spectrum of the voxels in the cell. We then calculated the median of these spectra across all cells in the tiles in the tile scan. We used this global median spectrum as an estimate of the fluorescent background. For each cell, we scaled the median spectrum such that the sum of values was equal to the sum of values from the cell’s spectrum, then we subtracted this scaled background estimate from the cell’s spectrum and zeroed negative values. We measured the cosine distance between each cell’s background subtracted spectrum and a set of reference spectra collected previously.^13^ For each cell, we identified the reference spectrum with the shortest cosine distance to the cell’s spectrum and mapped the resulting barcode assignment to a genus label.

### Cell density power spectrum

Given a set of cell locations for a genus, we constructed a cell density image by calculating the cell density in 10μm-by-10μm boxes separated by 5μm in the vertical and horizontal directions. Thus, the resolution for the density image is 5μm per pixel, and each pixel value is the density of a 100μm^2^ window. We then used PowerSpectrum tool from the Turbustat package with default parameters to calculate the power spectrum of the density image and perform linear regression.^32^

### Simpson diversity-area relationship

Given a set of cell locations and genus labels, we divided the tile scan image into a set of rectangle regions *A*_*j*_, *j* = 0,1, …, *n*_*A*_ where *n*_*A*_ is the total number of regions and each *A*_*j*_ has area *A*. We calculated the Simpson diversity *D* of each region,

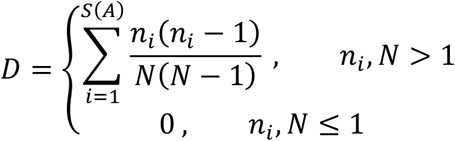

and calculated the sum of values for all regions,

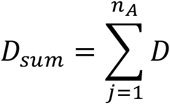

where *S*(*A*) is the number of genera in area *A, n*_*i*_ is the number of cells of genus *i* in area *A*, and *N* is the total number of cells in area *A*. We repeated this, dividing the tile scan into smaller rectangle regions with each repetition. We then fit the relationship between *D*_*sum*_ and *A* to a power law using linear regression on a log-log scale,

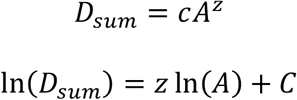

### Absolute abundance analysis

To calculate the volume of plaque imaged in a tile scan, we created a foreground mask to define the number of voxels with cells and multiplied by the voxel volume of the confocal point scan. For each genus, we divided the number of cells by the total volume imaged.

### Statistical analysis

No statistical methods are used to predetermine sample size. Unless otherwise stated, the experiments are not randomized. The investigators were not blinded to allocation during experiments and outcome assessment. Each data point in **Fig. 3b**,**c** represents the mean value of all tile scans of plaque biofilm from a given donor, including samples from different tooth aspects within the same patient. This donor grouping of tile scans is indicated by the “Donor ID” column in **Extended Data Table 5**. Boxplots consist of a bottom line representing the lower quartile (Q1), a line inside the box representing the median (Q2), a top line representing the upper quartile (Q3), an upper whisker extending from the top of the box indicating the maximum value within 1.5 times the interquartile range (IQR) above Q3 and a lower whisker extending from the bottom of the box indicating the minimum value within 1.5 times the IQR below Q1. For comparison across all groups, we used a Kruskal-Wallis H-test, then a post hoc two-tailed pairwise Dunn test. Significance levels were set at p ≤ 0.05 and p ≤ 0.01.

## Data Availability

Microscopy data have been deposited to Zenodo at https://doi.org/10.5281/zenodo.12659482,^41^ https://doi.org/10.5281/zenodo.12659548,^42^ https://doi.org/10.5281/zenodo.12659608,^43^ https://doi.org/10.5281/zenodo.12659599,^44^ https://doi.org/10.5281/zenodo.12659419,^45^ https://doi.org/10.5281/zenodo.12554811,^46^ https://doi.org/10.5281/zenodo.12630306,^47^ and https://doi.org/10.5281/zenodo.11493587.^48^

## Code Availability

The implementation of code to process data is available at https://github.com/benjamingrodner/peri_implantitis_HiPRFISH_analysis (v1.0.0).^40^

## Acknowledgements

We thank R. M. Williams and J. M. Dela Cruz for assistance with microscopy and Ioannis Ntekas and M. and H. Takayasu for discussions and feedback. This work was supported by the 2022 Osseointegration Foundation Basic Science Research Grant to D.M.K. and I.D.V., and by an instrumentation grant from the Kavli Institute at Cornell, by US National Institutes of Health (NIH) grants 1DP2AI138242 to I.D.V. and 1R33CA235302 to I.D.V., W.R.Z. and I.L.B. Imaging data were acquired in the Cornell Biotechnology Resource Center Imaging Facility using the shared, NYSTEM (CO29155)- and NIH (S10OD018516)-funded Zeiss LSM880 confocal and multiphoton microscope.

## Author Contributions

B.G., D.T.W., C.Y.C., D.M.K., and I.D.V. contributed to conception, design, and interpretation; B.G. and L.T. contributed to analysis; B.G., D.T.W., S.H., and N.W. contributed to data acquisition; B.G., D.T.W., C.Y.C., D.M.K., and I.D.V. drafted and critically revised the manuscript; All authors gave final approval and agreed to be accountable for all aspects of the work ensuring integrity and accuracy.

