## Extended Data for "Microscale Spatial Dysbiosis in Oral biofilms Associated with Disease"

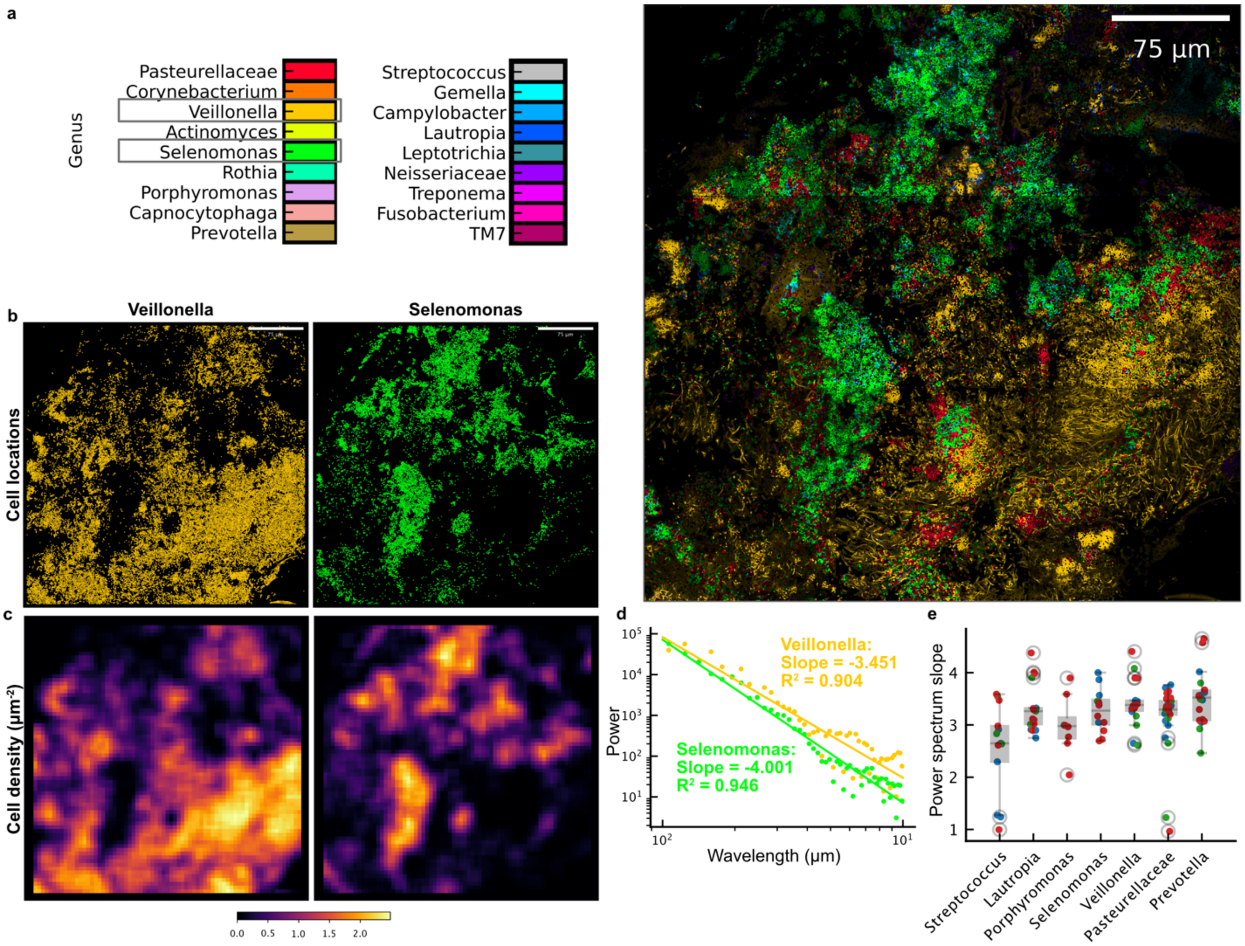


**Extended Data Figure 1: a** *Right:* Example processed image of healthy implant biofilm. *Left:* Genus color legend for the image. **b** Scatter plots of the locations of *Veillonella* and *Selenomonas* cells from the image in **a**. **c** Density maps of the *Veillonella* and *Selenomonas* cells. **d** Regressions of power spectra for the density of *Veillonella* and *Selenomonas* cells. Power spectrum values are calculated by taking radial profile of the square of the 2-dimensinal Fourier transform on the density map. **e** Summary of power spectrum slopes across all images, grouped by genus and colored by clinical status.

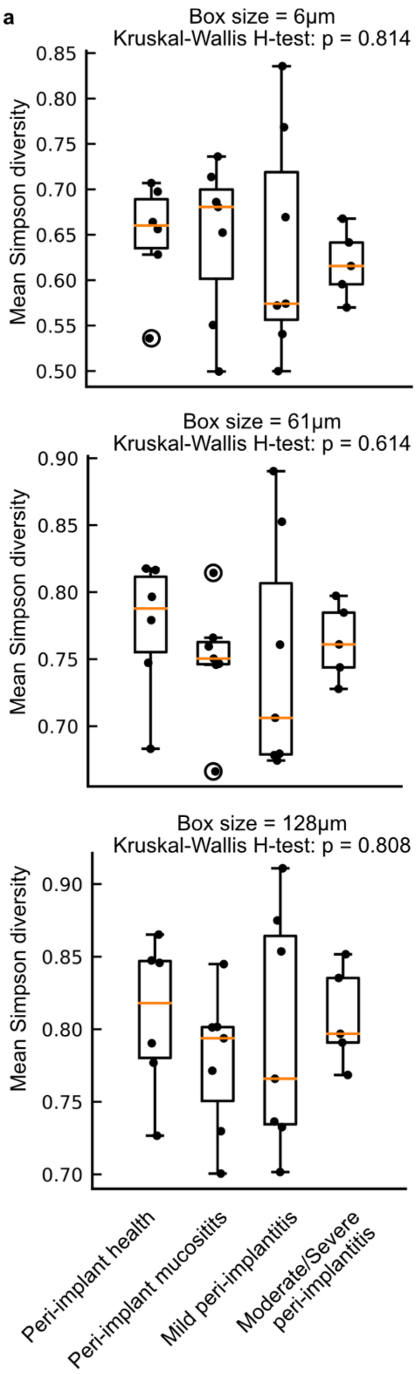


**Extended Data Figure 2: a** Mean Simpson diversity for all tile scans of each donor grouped by clinical status for a range of length scales. Tile scans were divided into boxes as in **Fig. 3a** and the Simpson diversity was calculated for each box, then the mean Simpson diversity was calculated across all tile scans for a donor. Kruskal-Wallis H-test indicated that we could not reject the null hypothesis that the population median of all groups are equal.

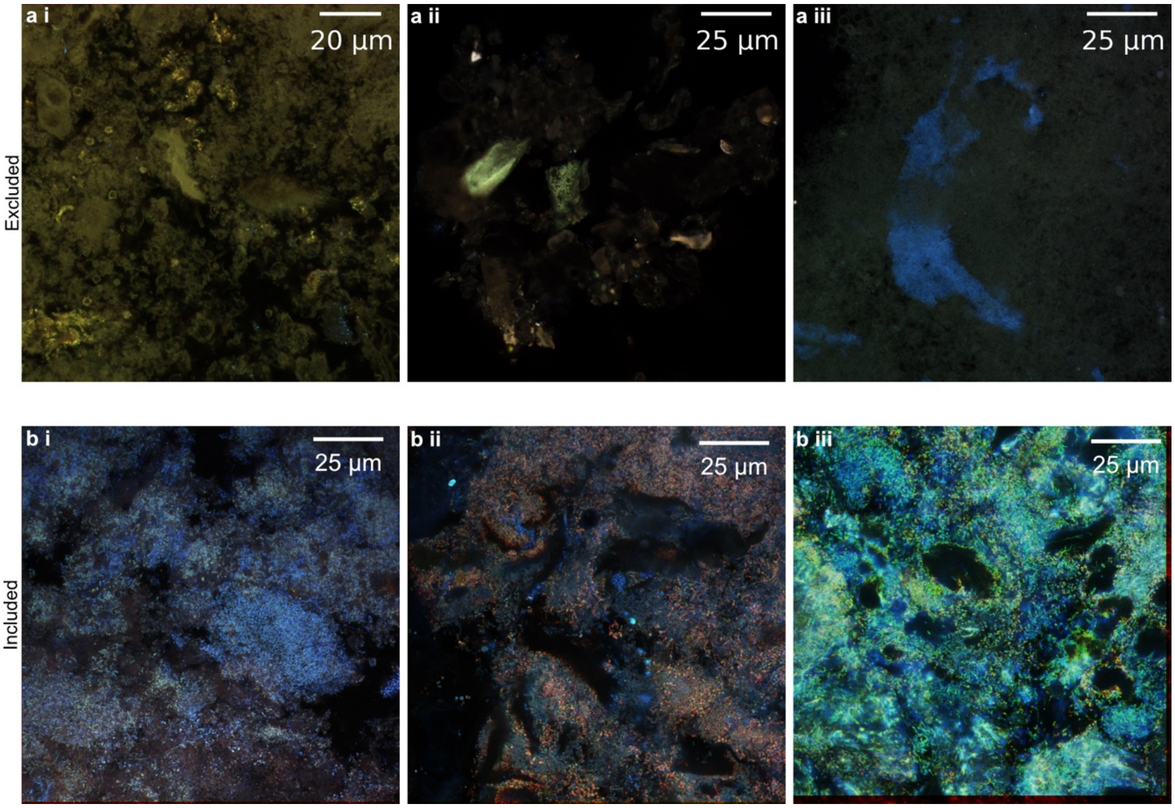


**Extended Data Figure 3:** Examples of excluded (**a**) versus included (**b**) images shown as RGB projections. To generate this RGB projection, spectral images from the 488nm, 514nm, 561nm lasers were summed along the spectral dimension and then normalized between 0 and 1 and stacked into an (x,y,3) array.
